# Aberrant peripheral immune responses in acute Kawasaki disease with single-cell sequencing

**DOI:** 10.1101/2020.11.05.369348

**Authors:** Zhen Wang, Lijian Xie, Sirui Song, Liqin Chen, Guang Li, Jia Liu, Tingting Xiao, Hong Zhang, Yujuan Huang, Guohui Ding, Yixue Li, Min Huang

## Abstract

Kawasaki disease (KD) is the most common cause of acquired heart disease in children in developed countries. Although diverse immune aberrance was reported, a global understanding of immune responses underlying acute KD was lacking. Based on single-cell sequencing, we profiled peripheral blood mononuclear cells from patients with acute KD before and after intravenous immunoglobulin therapy and from healthy controls. Most differentially expressed genes were derived from monocytes, with upregulation of immunoglobulin receptors, complement and receptors and downregulation of MHC class II receptors before therapy. The percentage of B cells was significantly increased before therapy and rapidly returned to normal after therapy. There was also an increased abundance of B-cell receptors with *IGHA* and *IGHG* after therapy, accompanied by massive oligoclonal expansion. The percentage of CD8 T cells was remarkably decreased during acute KD, especially the subset of effector memory CD8 T cells. All lymphocyte compartments were characterized by underexpressed interferon response pathways before therapy. The identification of unique innate and adaptive immune responses suggests potential mechanisms underlying pathogenesis and progression of KD.

## Introduction

Kawasaki disease (KD) is an acute, systemic febrile illness and vasculitis of childhood that can lead to coronary artery lesions (CALs)^1^. This disease predominantly affects children who are younger than 5 years and has become the most common cause of acquired heart disease among children in many developed countries^2^ The etiology of KD remains elusive, and diagnosis continues to depend on principal clinical features including fever, rash, conjunctivitis, changes in the oral mucosa and in the extremities, and cervical lymphadenopathy^3^. Because the signs and symptoms of KD resemble those of other childhood febrile illnesses, prompt diagnosis of KD is still challenging. High-dose intravenous immunoglobulin (IVIG) within the first 10 days after fever onset is the standard therapy for KD, which remarkably reduces the rate of CALs^4^ However, the mechanism of IVIG in the treatment of KD is unknown. Approximately 10% to 20% of KDs are IVIG resistant and are at increased risk for CALs^3^.

The most widely favored theory of KD etiology is that it is caused by a ubiquitous infectious agent in a small subset of genetically predisposed children^5^. Genome-wide wide association studies (GWAS) have identified a number of susceptibility genes for KD^6^. Many of the susceptibility genes including *ITPKC, CASP3, FCGR2A, CD40* and MHC class II are related to immune functions. Nonetheless, the causative agents initiating the disease have been still widely debated over 50 years. Various pathogens have been proposed as the trigger including superantigen toxin, Epstein-Barr virus, coronavirus and retrovirus, but none has been confirmed by subsequent studies^7^ Despite the uncertainty of the cause, activation of the immune system provides important evidence for its pathogenesis. Plasma levels of pro-inflammatory cytokines such as TNF-α, IL-1β, and IFN-γ are elevated during the acute phase of KD^8^. Innate immune cells including neutrophils and monocytes are elevated in the peripheral blood and have been identified in the arterial wall early in the disease^9^ Activation of peripheral blood lymphocytes seems also responsible for the development of KD, though their roles remain controversial^10^. In addition, antigen-specific IgA plasma cells and CD8 T cells infiltrate inflamed tissues, implying an immune response to an intracellular pathogen entering through the respiratory tract^11^. Recently, a high incidence of Kawasaki-like disease has been reported during the severe acute respiratory syndrome coronavirus 2 (SARS-CoV-2) epidemic^12,13^, suggesting a possible link between coronavirus infection and KD.

The immune system is composed of numerous cell types with varying states. Most of previous studies performed immunophenotyping based on flow cytometry, which is constrained to a few selected cell types and markers. Several studies performed transcriptome analyses of KD on bulk cell populations^14,15^, but the heterogeneity of the immune system cannot be resolved. Over the past few years, the revolution in single-cell sequencing has enabled an unbiased quantification of gene expression in thousands of individual cells, which provides an more efficient tool to decipher the immune system in human diseases^16^. In this study, we profiled peripheral blood mononuclear cells (PBMCs) isolated from acute KDs at single-cell resolution. Our results revealed global and dynamic immune responses unique to each cell compartment before and after IVIG therapy.

## Results

### Study design and single-cell profiling of PBMCs

We collected eight fresh peripheral blood samples derived from four patients with acute KD (P1-P4, Supplementary Table S1). The patients were diagnosed according to the criteria proposed by the American Heart Association^3^. Three patients met criteria for complete KD and one for incomplete KD. For each patient, the first blood sample was taken on the 5th days after the onset of fever before IVIG therapy. The second sample was obtained at 24 hours after completion of IVIG therapy and subsidence of fever (Supplementary Table S2). All patients were IVIG sensitive responders and had not developed CALs. We also collected fresh peripheral blood samples from three age-matched healthy donors as controls (H1-H3, Supplementary Table S1).

We used the 10× Genomics platform for single-cell RNA sequencing (scRNA-seq) of PBMCs isolated from the samples. The total number of detected cells passing quality control were 57,881, including 19,319 cells for patients before IVIG therapy, 23,874 cells after therapy, and 14,688 cells for healthy controls (Supplementary Table S3). We also performed single-cell B cell receptor sequencing (scBCR-seq) and single-cell T cell receptor sequencing (scTCR-seq) based on the scRNA-seq libraries. Totally, 15,419 and 40,841 cells with productive paired BCRs and TCRs were detected, respectively (Supplementary Table S4 and Table S5).

Based on the scRNA-seq profiles, we clustered the cells across samples and visualized them in two-dimensional space (Figure 1A). Major cell compartments comprising PBMCs could be characterized by canonical marker genes (Figure 1B), including T cells *(CD3D, CD3E, CD3G,* 60.20%), CD4 T cells (*CD4*, 38.95%), CD8 T cells *(CD8A, CD8B,* 21.25%), natural killer (NK) cells *(NCAM1* or *CD56, KLRB1, NKG7,* 7.35%), B cells *(CD19, MS4A1* or *CD20, CD38,* 20.96%), monocytes *(CD14, CD68, FCGR3A* or *CD16,* 7.55%) and plasmacytoid dendritic cells (pDCs) *(LILRA4,* 0.23%). There were also some residual erythrocytes (*HBB*, 2.49%) and megakaryocytes (*PPBP*, 0.83%) mixed in the PBMCs, which were removed in the following analyses.

**Figure 1.**
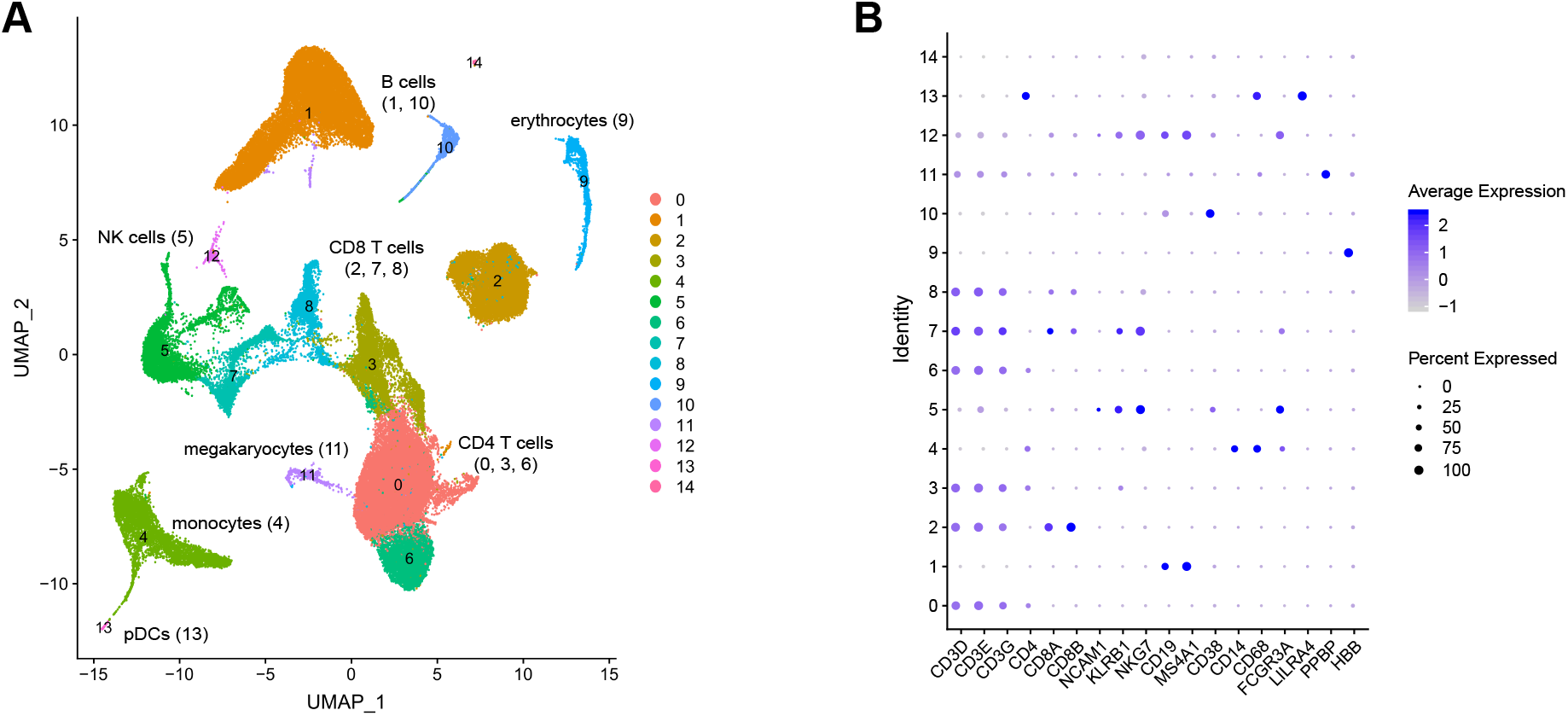
Major cell compartments of PBMCs revealed by scRNA-seq. (A) UMAP visualization of cells, where cell identity are indicated by cluster numbers and colors. Cell types were annotated based on the expression of canonical marker genes. (B) Dot plot depicting average expression and percentage expression of canonical marker genes in each cell cluster.

### Overall dynamics of cell abundance

We compared the percentage of each cell compartment between KDs before and after IVIG therapy, as well as healthy controls (Figure 2A). To verify the scRNA-seq results, we also retrieved a clinical dataset of large sample size, which comprised flow-cytometric estimation of peripheral blood lymphocytes from 125 acute KDs before IVIG therapy (Figure 2B). The scRNA-seq data demonstrated that the B cell abundance increased significantly in KDs before therapy, compared with those after therapy *(P* = 0.01) and in healthy controls *(P* = 0.04, two-sided t-test, Figure 2A). The increase of B cells in acute KD was observed in several previous studies^10,17–19^, and our flow-cytometric dataset also confirmed a significantly higher levels of B cells than the reference range in both percentages (*P* = 4.74×10^-20^, Figure 2B) and absolute counts (*P* = 1.59×10^-7^, two-sided t-test, Supplementary Figure S1). The scRNA-seq data also showed a significantly lower abundance of CD8 T cells in KDs than healthy controls (*P* = 0.02, two-sided t-test), though no significant difference were found before and after IVIG therapy (Figure 2A). The decrease of CD8 T cells was consistent with previous reports^10,17–19^, which could be confirmed by our flow-cytometric dataset in both percentages (*P* = 2.84×10^-34^, Figure 2B) and absolute counts (*P* = 5.00×10^-13^, two-sided t-test, Supplementary Figure S1).

**Figure 2.**
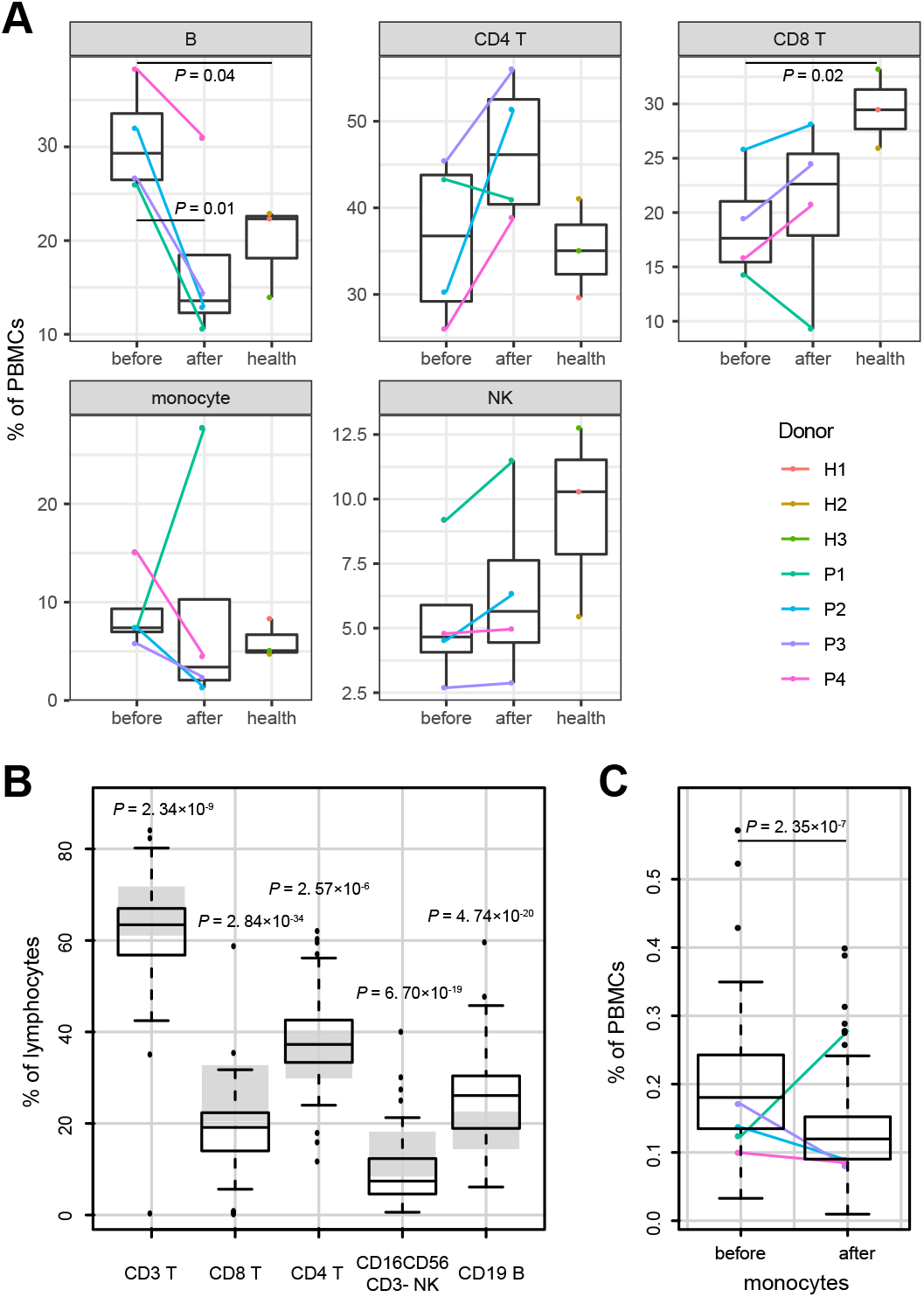
Box plots for comparison of cell abundance. (A) Percentage of each cell compartment in PBMCs between pre- and post-therapy patients and healthy controls. (B) Percentage of lymphocyte compartments by flow cytometry from 125 KDs before IVIG therapy. The gray area represents the reference range. (C) Percentage of monocytes by routine blood test from 80 KDs before and after IVIG therapy. Samples used for scRNA-seq are highlighted. Box plots represent the interquartile range (IQR), with horizontal lines indicating the median value, and whiskers extending to the farthest data point within a maximum of 1.5 × IQR. All *P*-values are calculated using two-sided t-tests.

A reduction of NK cells was reported in acute KD^18,20,21^ and can be verified by our flow-cytometric dataset (*P* = 6.70×10^-19^, two-sided t-test, Figure 2B and Supplementary Figure S1). The tendency of NK cell alternations was consistent with the scRNA-seq data, though statistical significance was lacking (Figure 2A). There remained a controversy in literature regarding whether the level of CD4 T cells was increased or decreased in acute KD^10,17–19^. While our flow-cytometric dataset showed a significant increase of CD4 T cells in percentages of lymphocytes (*P* = 2.57×10^-6^, two-sided t-test, Figure 2B), they did not differed significantly in absolute counts compared with the reference range (Supplementary Figure S1). We did not find clear tendency of CD4 T cell alternations in the scRNA-seq data (Figure 2A).

It was reported that the abundance of monocytes was decreased following IVIG therapy^10,22,23^. However, one patient (P1) in the scRNA-seq data showed remarkable increase of monocytes after therapy (Figure 2A). To exclude any technical bias, we retrieved the routine blood test dataset of the patients, as well as other 80 acute KDs before and after IVIG therapy (Figure 2C and Supplementary Figure S2). The dataset confirmed a significant decrease of monocytes after therapy in general (*P* = 2.35×10^-7^, two-sided t-test), whereas P1 displayed a reverse tendency as in the scRNA-seq data.

### Monocyte subsets and responses

We further re-clustered the monocytes into three distinct subsets to dissect their heterogeneity (Figure 3A and Supplementary Figure S3), including CD14 monocytes *(CD14,* 54.78%), CD16 monocytes *(FCGR3A,* 38.37%) and myeloid dendritic cells (mDCs) *(CD1C,* 6.85%). After IVIG therapy, the patients showed a substantially decreased abundance of CD16 monocytes (*P* = 3.02×10^-3^, two-sided t-test, Figure 3B). It was reported that CD16 monocytes were more closely associated with many inflammatory conditions^24^, including acute KD^10,23^. The abundance of CD16 monocytes remained significantly lower in patients after therapy than healthy controls (*P* = 0.04, two-sided t-test), but we did not find significant difference in patients before therapy and healthy controls (Figure 3B).

**Figure 3.**
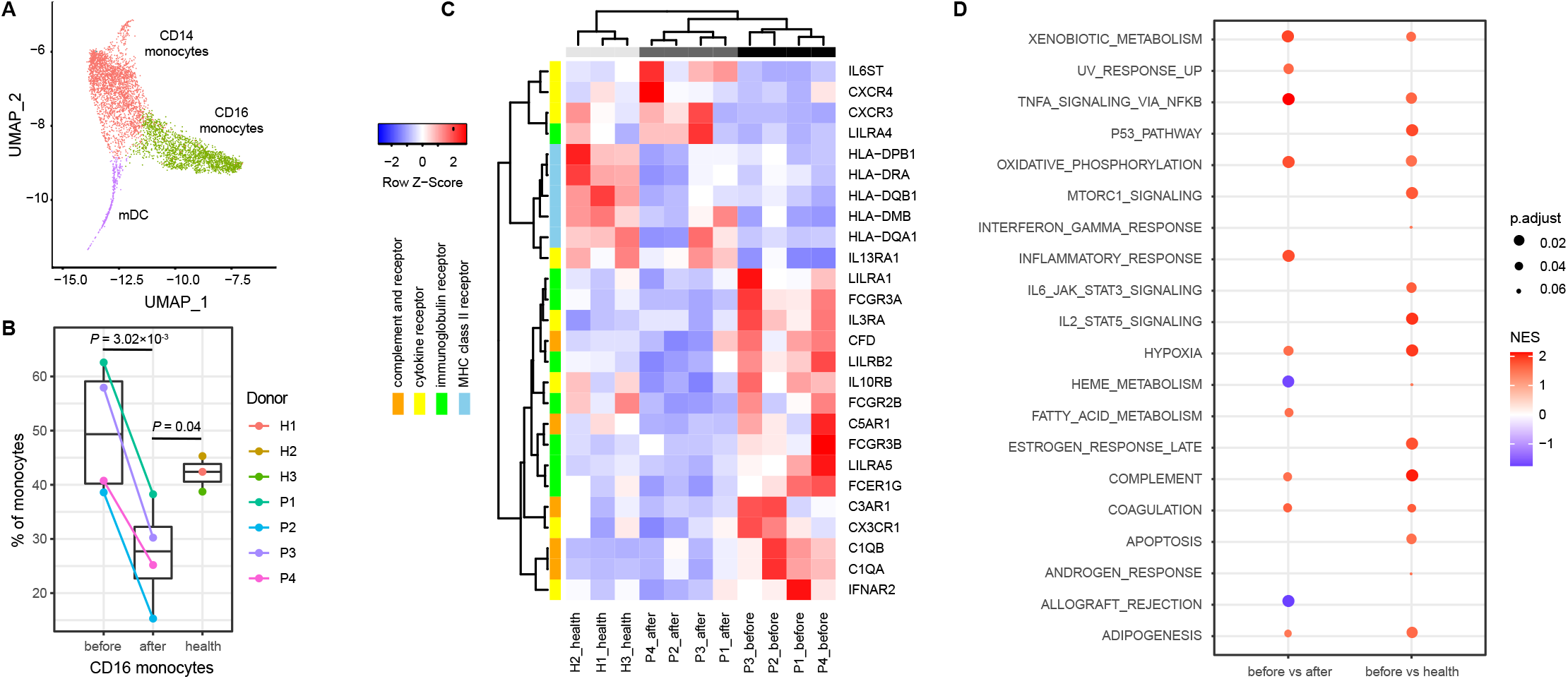
Characterization of monocytes and transcriptional signatures. (A) UMAP plot of monocytes colored by their subsets. (B) Percentage of CD16 monocytes across conditions. *P*-values are calculated using two-sided t-tests. (C) Heat map of DEGs with functional enrichment. Gene expression on the sample level is measured by the counts per million, which is then normalized across samples by the Z-score. (D) Dot plot depicting significant hallmark gene sets by GSEA. The dot size indicates the adjusted *P*-value (FDR), and the dot color indicates the normalized enrichment score (NES). Pre-therapy samples are compared with post-therapy ones and healthy controls, respectively. A positive NES suggests that the gene set is enriched in upregulated genes before therapy, and a negative NES suggests it is enriched in downregulated genes before therapy.

To identify more genes involved in the immune responses, we performed differential expression analysis for each cell type. The most differentially expressed genes (DEGs, false discovery rate [FDR] < 0.1) resulted from monocytes among all PBMC compartments (Supplementary Figure S4). Gene ontology (GO) and KEGG pathway^25^ analyses for the DEGs suggested that they were significantly enriched in immunoglobulin receptors, cytokine receptors, complement and receptors and MHC class II receptors (FDR <0.1, Supplementary Figure S5). Most DEGs encoding immunoglobulin receptors were upregulated in patients before IVIG therapy compared with those after therapy and healthy controls, including *FCGR3A*, *FCGR3B*, *FCGR2B* and *FCEG1G* (Figure 3C). The reduced expression of *FCGR3A* and *FCGR2B* after IVIG therapy in monocytes were in agreement with previous studies^26,27^. Most DEGs of complement and receptors were also upregulated before therapy, including *C1QA*, *C1QB*, *CFD*, *C3AR1* and *C5AR1* (Figure 3C). In contrast, most DEGs encoding MHC class II receptors were downregulated in acute KD than healthy controls regardless of therapy, including *HLA-DQA1*, *HLA-DQB1*, *HLA-DRA*, *HLA-DPB1* and *HLA-DMB* (Figure 3C). The expression patterns of DEGs encoding cytokine and chemokine receptors were more diverse. While some were upregulated in patients before therapy including *IFNAR2, IL3RA, IL10RB* and *CX3CR1,* others were overexpressed after therapy or in healthy controls including *IL13RA1, IL6ST, CXCR3* and *CXCR4* (Figure 3C).

We then performed the gene set enrichment analysis (GSEA)^28^ to detect changes of pathway activity, which did not depend on predefined DEGs but coordinated changes of functionally related genes. In comparison with patients after therapy or healthy controls, a large number of pathways related to cytokine signaling and inflammatory responses were upregulated before therapy (FDR < 0.1, Figure 3D). Especially, TNF signaling via NF-kB, complement and coagulation pathway were activated in both comparisons. The activation of NF-kB in monocytes of acute KD was previously reported^29^.

### B cell subsets and BCR repertoires

We re-clustered B cells and identified three distinct B cell subsets (Figure 4A and Supplementary Figure S6): naive B cells *(MS4A1, TCL1A,* 67.20%), memory B cells *(MS4A1, CD27, IGHA1, IGHG1,* 26.34%) and plasma cells *(CD27, CD38, IGHA1, IGHG1,* 6.46%)^30^. After ΓVIG therapy, the proportion of plasma cells were increased compared with that before therapy, although the result just reach the statistical significance level (*P* = 0.05, two-sided t-test, Figure 4B). As few DEGs could be detected in B cells, we performed GSEA to assess the pathway activity (FDR < 0.1, Figure 4C). We found that interferon response pathways were underexpressed before IVIG therapy compared with those after IVIG therapy and in healthy controls. In addition, many pathways related to inflammatory responses were upregulated after therapy, which was in contrast to the cases in monocytes. Pathways associated with cell cycles were upregulated before therapy including G2M checkpoint and E2F targets (Figure 4C), which might be responsible for the high abundance of B cells in acute KD.

**Figure 4.**
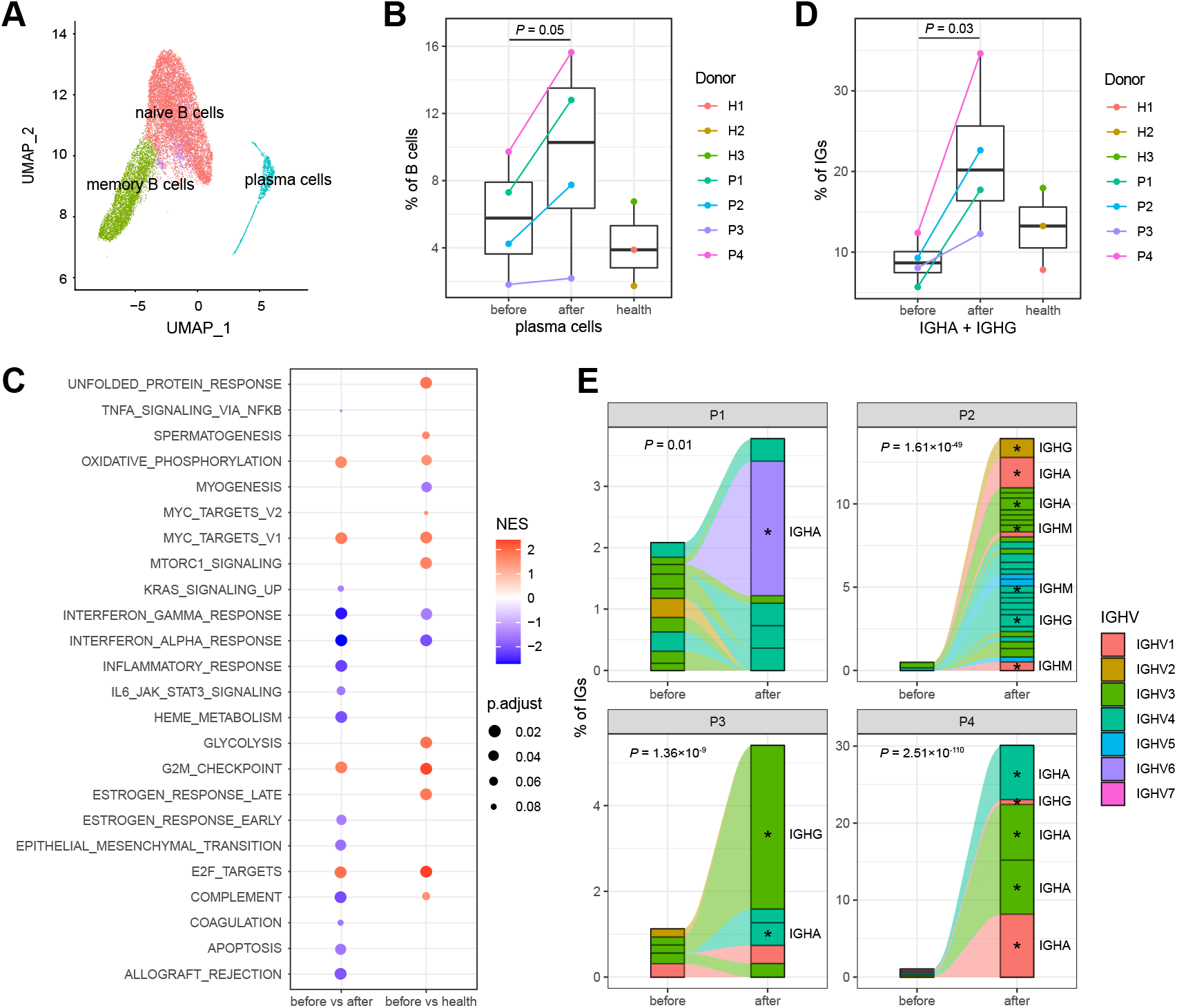
Characterization of B cells and BCRs. (A) UMAP plot of B cells colored by their subsets. (B) Percentage of plasma cells across conditions. *P*-values are calculated using two-sided t-tests. (C) Significant hallmark gene sets by GSEA. The dot size indicates the adjusted *P*-value (FDR), and the dot color indicates the normalized enrichment score (NES). A positive NES suggests that the gene set is enriched in upregulated genes before therapy, and vice versa. (D) Percentage of *IGHA* and *IGHG* in BCRs across conditions. *P*-values are calculated using two-sided t-tests. (E) BCR clonotype tracking in each patient. The clonotype composition is represented by stacked bar plots, which are colored according to *IGHV* families. Only clonotypes with clonal size ≥ 3 are plot, and proportion of BCRs in these clonotypes are compared before and after therapy by using Fisher’s exact tests. Individual clonotypes with FDR < 0.05 are marked with stars.

We analyzed the scBCR-seq data to explore dynamics of BCR repertoires during acute KD. The abundance of *IGHA* and *IGHG* were significantly elevated after IVIG therapy (*P* = 0.03, two-sided t-test, Figure 4D and Supplementary Figure S7), which implicated isotype switching of BCRs from IgM/IgD to IgG/IgA resulted from B cell activation. We then examined the clonality of BCRs based on unique VDJ sequences for each patients. In comparison with patients before therapy, the percentage of clonal BCRs (clonal size ≥ 3) was greatly increased after therapy (Figure 4E). Particularly, in every patient, we could observe remarkable oligoclonal expansions after therapy. The significantly expanded clones were dominated by the isotype IgA and IgG (FDR < 0.05, Fisher’s exact test, Figure 4E), suggesting that antibody-mediated responses might be involved in the recovery of acute KD. The *IGHV* gene usage of the expanded clones was diverse among the patients (Figure 4E), indicating specific B cell reactions rather than responses to a superantigen. We also did not find obvious bias in *IGHV* usage across patients and healthy controls (Supplementary Figure S8). Oligoclonal IgA plasma cells infiltrate vascular tissue in acute KD were reported in previous studies^31,32^, and synthetic antibodies were designed to detect pathogens of KD^33,34^ Our analyses suggested that the antibodies might also be identified from peripheral blood of patients recovered from acute KD.

### T and NK cell subsets and TCR repertoires

We subdivided CD4 T, CD8 T and NK cells based on expression of canonical genes, respectively (Figure 5A). Both CD4 T and CD8 T cells comprised three subsets (Supplementary Figure S9): naive cells (*CCR7*, *SELL,* 70.67%), central memory cells (co-expression of *CCR7, SELL* and *S100A4,* 13.12%) and effector memory cells *(S100A4,* 10.82%)^35^. The memory CD8 T cells also expressed high levels of cytotoxicity-associated genes *(GZMK, GZMB).* In addition, regulatory CD4 T cells *(FOXP3,* 3.45%) and proliferating T cells *(TYMS,* 1.95%) were identified. While we did not observe significant alterations among the CD4 T cell subsets during acute KD (Supplementary Figure S10), we found a significantly decreased percentage of effector memory cells (*P* = 0.01) and increased percentage of naive cells in the CD8 T cells (*P* = 0.002, two-sided t-test, Figure 5B). These results indicated that the effector memory cells, which can acquire rapid cytotoxic functions following activation, were the major subset contributing to the reduction of CD8 T cells in PBMCs. Interestingly, CD8 T cells were reported to infiltrate coronary arteries in acute KD^36^, suggesting that the reduction of CD8 T cells in the peripheral blood might be involved in the clearance of pathogens. The NK cells included two subsets (Supplementary Figure S11): CD16-high cells *(FCGR3A,* 88.63%) and CD56-high cells *(NCAM1, KLRC1,* 11.37%)^37^. Although we observed a tendency of decreased abundance of CD16-high cells as in monocytes, the result was not statistically significant (Supplementary Figure S12).

**Figure 5.**
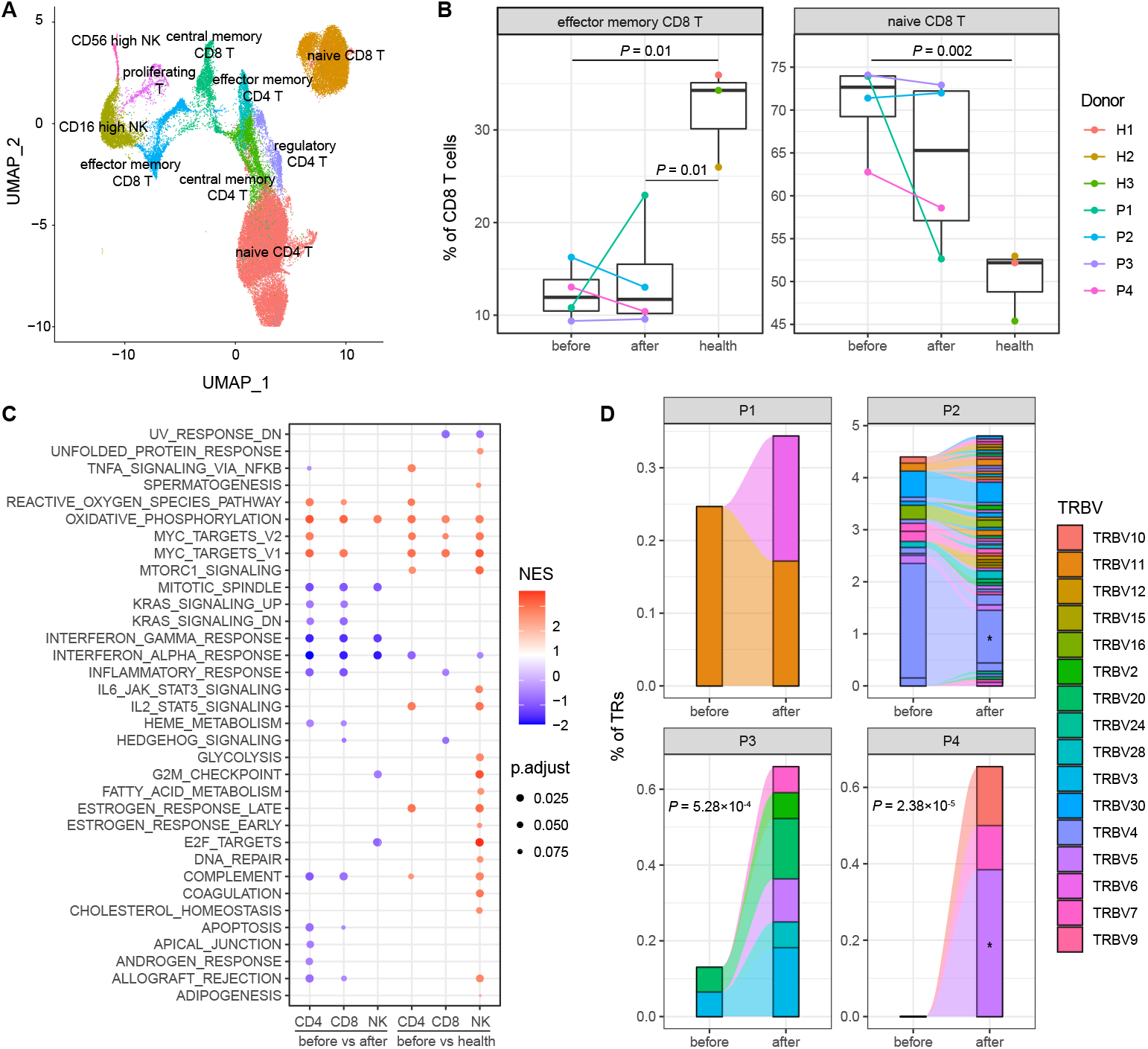
Characterization of T and NK cells and TCRs. (A) UMAP plot of T and NK cells colored by their subsets. (B) Percentage of effector memory and naïve CD8 T cells across conditions. *P*-values are calculated using two-sided t-tests. (C) Significant hallmark gene sets by GSEA. The dot size indicates the adjusted *P*-value (FDR), and the dot color indicates the normalized enrichment score (NES). A positive NES suggests that the gene set is enriched in upregulated genes before therapy, and vice versa. (D) TCR clonotype tracking in each patient. The clonotype composition is represented by stacked bar plots, which are colored according to *TRBV* families. Only clonotypes with clonal size ≥ 3 are plot, and proportion of TCRs in these clonotypes are compared before and after therapy by using Fisher’s exact tests. Individual clonotypes with FDR < 0.05 are marked with stars.

We profiled pathway activity by GSEA for CD4 T, CD8 T and NK cells (FDR < 0.1). Many common pathways were associated with these lymphocyte compartments, such as overexpression of oxidative phosphorylation and Myc targets before therapy (Figure 5C). Notably, all of the lymphocyte compartments showed lower activity of interferon response pathways in patients before therapy than those after therapy or healthy controls (Figure 5C). These results were similar to B cells and suggested that deficiency in interferon responses of lymphocytes could be important in the pathogenesis of acute KD. In contrast to T cells, NK cells displayed extensive activation of many cytokine signaling and complement pathways in pre-therapy patients compared with healthy controls, which might result from their innate immune responses as monocytes.

Finally, we tracked the TCR clonotypes based on the scTCR-seq data. T cells with clonal TCRs (clonal size ≥ 3) were enriched in effector memory CD8 T cells, especially those with high expression of *GZMB* (Supplementary Figure S13), indicating strong cytolytic activity of the cells. We observed the percentage of clonal TCRs was significantly increased only in two patients (P3 and P4) after IVIG therapy (Figure 5D). A possible explanation was that the clonal expansion of TCRs was initiated before therapy. We did not find any predominant *TRBV* families in the clonal TCRs (Figure 5D), and no skewed usage of *TRBV* and *TRAV* could be detected by comparing patients and healthy controls (Supplementary Figure S14 and S15). These results suggested that conventional antigens^38–40^ rather than a superantigen^41–44^ were involved in the pathogenesis of KD.

## Discussion

Although great efforts have been made in the research and treatment of KD, many essential questions remain to be clarified, such as what trigger the disease, how the CAL is developed and the mechanism of IVIG therapy. The lack of knowledge has impeded a precise diagnostic test, a better management of IVIG resistant patients and long-term cardiac sequelae. An overt immune reaction is the most important clinical features of acute KD, which has helped to elucidate etiology and pathogenesis of the disease^5,7^ However, many immunological studies reported conflicting results^10^, in part due to the limited resolution of flow cytometry. Transcriptome-level analyses were performed for peripheral blood, but the signals were overwhelmed by the activation of innate immune system due to the dominate abundance of neutrophils^14,15^. In this study, we used single-cell sequencing to dissect the complex immune responses of PBMCs in acute KD. We observed remarkable temporal changes in the cell abundance before and after IVIG therapy, which were largely consistent with previous studies and our flow-cytometric dataset. In particular, the abundance of B cells, CD8 T cells and NK cells could be helpful in distinguishing KD and other febrile diseases, and predicting the effectiveness of IVIG^18^.

We identified unique signatures of gene expression for each cell compartment. Most of the DEGs arose from monocytes, in agreement with the immune dysregulation primarily occurring in the innate immune system^9^. Many DEGs encoding immunoglobulin receptors showed high expression level before IVIG therapy and returned to normal level after therapy, suggesting that the immunomodulatory effects of IVIG may involve the blockade of activating immunoglobulin receptors^45^. Another intriguing signature was the downregulation of MHC class II genes in acute KD relative to healthy controls, which was robustly observed in patients with SARS-CoV-2 infection^46^. Interestingly, the gene cluster of immunoglobulin receptors and MHC class II receptors were also identified as susceptibility loci of KD in GWAS studies^6^, highlighting their important roles in KD pathogenesis.

Regarding to the adaptive immune system, we found significant depletion of CD8 T cells in PBMCs during acute KD, especially the subset of effector memory cells. In addition, the subset of B cells expressing *IGHA* and *IGHG* showed elevated abundance after IVIG therapy, which was accompanied by oligoclonal expansion of BCRs. These results were consistent with an acquired immune response to specific antigens. In previous studies, the observation of CD8 T cells in inflamed tissues coupled with the presence of IgA plasma cells were proposed as the most important evidences for a respiratory viral etiology of KD^5^. Synthetic antibodies based on the oligoclonal IgAs were also designed to identify potential antigens^11^. However, clinical material available for the study is scarce because accessing the inflamed tissue of KD is impossible in living children. Our results suggested that the clonal BCRs prepared from peripheral blood of patients who recover from acute KD could show promising opportunity for antigen identification and therapeutic development. In contrast to innate immune cells such as monocytes and NK cells, our pathway analyses indicated that the inflammatory responses in T and B cells were much weaker in pre-therapy patients than healthy controls. After IVIG therapy, a common signal for all lymphocyte compartments was the activation of interferon response pathways. Several transcriptome analyses reported a striking absence of interferon-induced gene expression in patients with KD, especially compared with virus infection^15,47^. *IFNG* polymorphisms were also associated with the susceptibility of KD and IVIG responsiveness^48^. If KD was triggered by viral infection, it was plausible that the deficiency in interferon response pathways could be responsible for KD pathogenesis.

There were several shortcomings in our study. The sample size for single-cell analysis was limited, which affected the statistical power in differential abundance and differential expression analysis. Because all patients diagnosed with acute KD would be treated with IVIG, the intrinsic immune responses to potential pathogens were mixed with the immunomodulatory effects of IVIG. For example, further efforts should be made to distinguish clonally expanded BCRs specific to unknown pathogens and exogenous immunoglobulins after therapy. Also, more patients with various clinical presentation are needed to determine the relationship between immune responses of different cell types and disease outcomes.

## Materials and Methods

### Patients

All patients were recruited with informed consent from Shanghai Children’s Hospital between December 2019 and June 2020. The study was approved by the ethics committee of the Shanghai Children’s Hospital. The diagnosis of KD was made by using the criteria proposed by the American Heart Association^3^, including fever of unknown etiology lasting for ≥5 days, and the presence of five associated symptoms (conjunctivitis, oral changes, extremity changes, rash, cervical lymphadenopathy). Complete KD was diagnosed if at least four of the symptoms were fulfilled, otherwise incomplete KD. The patients received high-dose IVIG (1 g/kg per day) for two consecutive days combined with oral aspirin after diagnosed with KD. CALs were regularly monitored by two-dimensional echocardiography.

### Single-cell preparation and sequencing

For each donor, 2 ml venous blood was collected in EDTA tubes and transferred to the laboratory with ice. The blood was processed within 4 hours of collection. PBMCs were isolated by density gradient centrifugation using the Ficoll-Paque medium. The cell viability should exceed 80% determined with trypan blue staining. Single-cell capturing and library construction were performed using the Chromium Next GEM Single Cell V(D)J Reagent kits v1.1 (10x Genomics) according to the manufacturer’s instructions. The constructed library was sequenced on the Illumina NovaSeq platform.

### scRNA-Seq data analysis

The raw sequencing data were processed by the Cell Ranger pipeline (v3.0.1, 10x Genomics), including demultiplexing, genome alignment (GRCh38), barcode counting and unique molecular identifier (UMI) counting. The gene-barcode matrix of UMI counts was then analyzed with Seurat (v3.0.2)^49^ For most samples, the following criteria were applied for quality control: total UMI count between 2,000 and 60,000, and mitochondrial gene percentage < 5%. For P1 before therapy, a lower cutoff of total UMI count (1,000) was used due to its lower median UMI count per cell. The count matrix was log-normalized, and the top 5,000 most variable genes were identified for dimensional reduction. All samples were integrated to remove batch effects based on the first 20 dimensions of the canonical correlation analysis (CCA). The integrated matrix was then scaled, and the first 30 dimensions resulted from the principal component analysis (PCA) were used for the uniform manifold approximation and projection (UMAP) visualization. Meanwhile, graph-based clustering was performed on the PCA-reduced data to identify cell clusters. The resolution was set to 0.1 to obtain major compartments of PBMCs, and 1.2 to obtain subsets of each compartment. The cell clusters were manually annotated based on the expression of canonical marker genes.

### Differential expression and functional enrichment analysis

We performed differential expression analysis for each cell compartment on the sample level following the tutorial of Bioconductor^50^. A pseudo-bulk expression profile was generated by summing counts together for all cells with the same combination of compartment and sample. Then the differential expression analysis was conducted between conditions by using edgeR (v3.30.3)^51^, which modelled the overdispersed count data based on a negative binomial generalized linear model. We used clusterProfiler (v3.16.0)^52^ for functional enrichment analyses. The GO and KEGG over-representation analysis were performed for DEGs with FDR < 0.1. The GSEA was performed for gene lists sorted by the log(fold-change), and the hallmark gene sets in MSigDB (v7.1)^53^ were used for annotation. *P*-values were adjusted by FDR and a gene set was considered significant with FDR < 0.1.

### BCR and TCR data analysis

The scBCR-seq and scTCR-seq data were assembled by the Cell Ranger VDJ pipeline (v3.0.1, 10x Genomics), and reference sequences in IMGT^54^ were fetched for annotation. Only cells with productive and paired chains (IGH and IGL/IGK for BCRs, TRA and TRB for TCRs) were preserved. For cells with more than one consensus sequence detected for the same chain type, the one with higher UMIs was chosen. Clonotypes represented by CDR3 sequences and gene usages were explored by immunarch (v0.6.5)^55^. The abundance of a clonotype was compared between pre- and post-therapy samples by the Fisher’s exact test. A clonotype must occur ≥3 times in at least one sample to be considered. Clonotypes and expression profiles were matched based on the same cell barcode.

## Data Availability

The raw sequence data reported in this paper have been deposited in the National Omics Data Encyclopedia (https://www.biosino.org/node/) of the Bio-Med Big Data Center, Shanghai Institute of Nutrition and Health, Chinese Academy of Sciences, under accession code OEP001162.

## Acknowledgements

This work was supported by the National Key R&D Program of China (2016YFC0901704, 2017YFC1201200, 2017YFA0505500), the Chinese Academy of Sciences (KFJ-STS-QYZD-126), the Youth Innovation Promotion Association CAS (2017325), the Hospital Development Center research funding (SHDC12016119), the Shanghai Science and Technology Committee research funding (17411954300), the Shanghai Municipality Health Commission research funding (2019SY025), and the Shanghai Children’s Hospital research funding (2019YLYZ01). We would like to thank all the patients who contributed samples, Anhui Engineering Laboratory for Big Data of Precision Medicine who contributed computational resource, and Genergy Bio-technology, Co. Ltd who provided single-cell sequencing support for this work.

## Author Contributions

Z.W., L.X., G.D., Y.L. and M.H. designed research. L.X., S.S., L.C. and M.H. collected samples. Z.W., G.L., J.L., G.D. and Y.L. analyzed data. T.X., H.Z. and Y.H. provided clinical information. Z.W., L.X., G.D., Y.L. and M.H. wrote the manuscript.

