## Supplementary for "Aberrant peripheral immune responses in acute Kawasaki disease with single-cell sequencing"

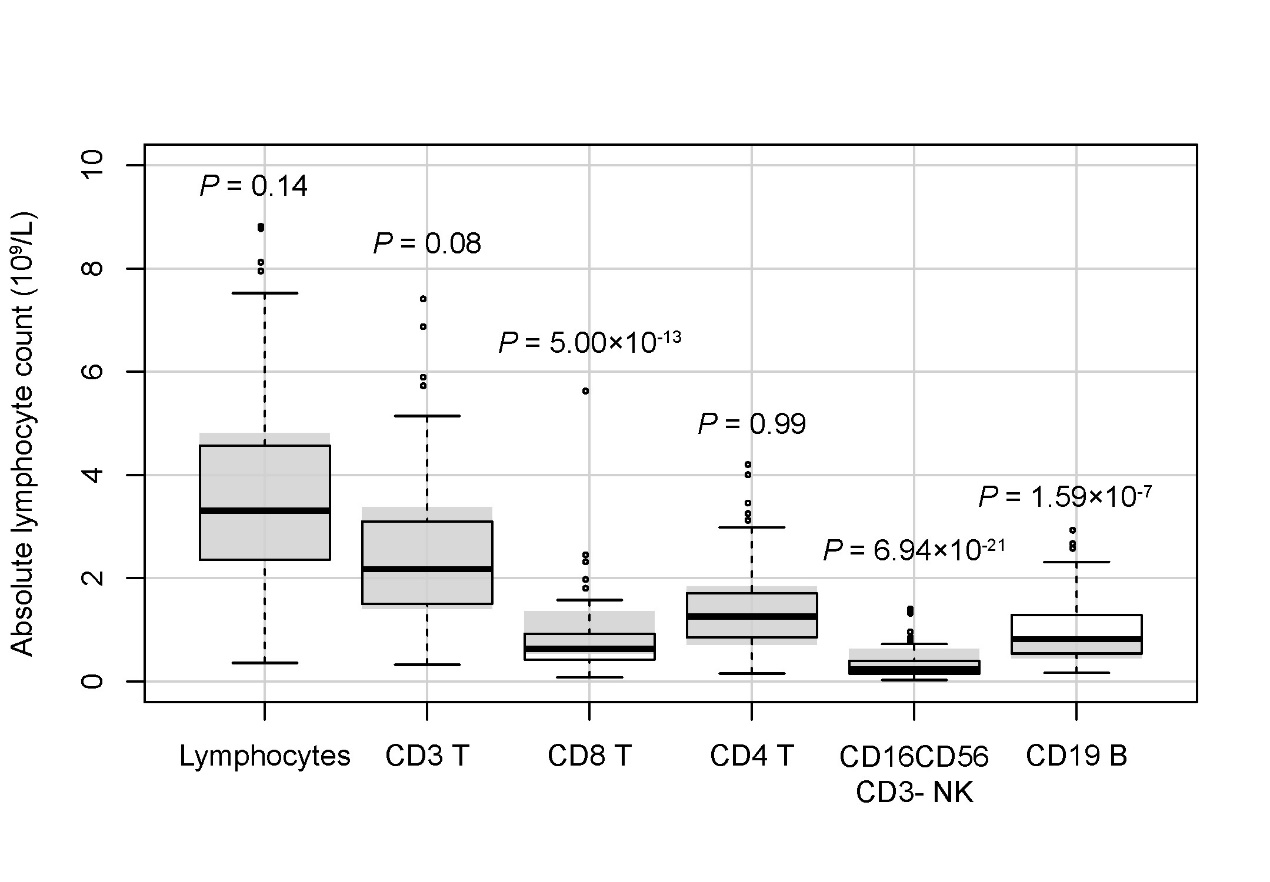

Supplementary Figure S1. Absolute lymphocyte count by flow cytometry for 125 acute KDs before IVIG therapy. The gray area represents the reference range. The P-values are calculated in comparison to the reference range using two-sided t-tests.

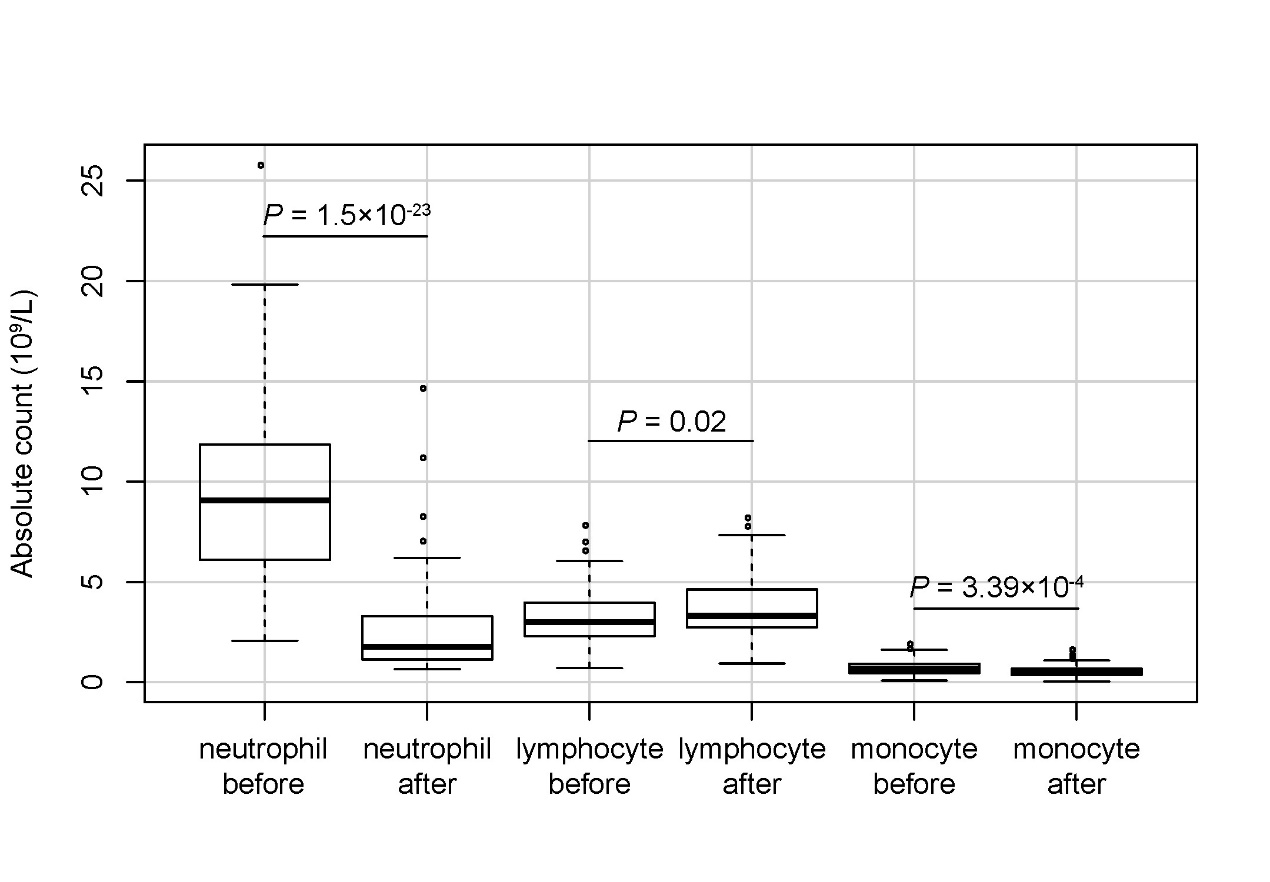

Supplementary Figure S2. Absolute cell count by routine blood test for 80 acute KDs before and after IVIG therapy. The P-values are calculated between pre- and post-therapy samples using two-sided t-tests.

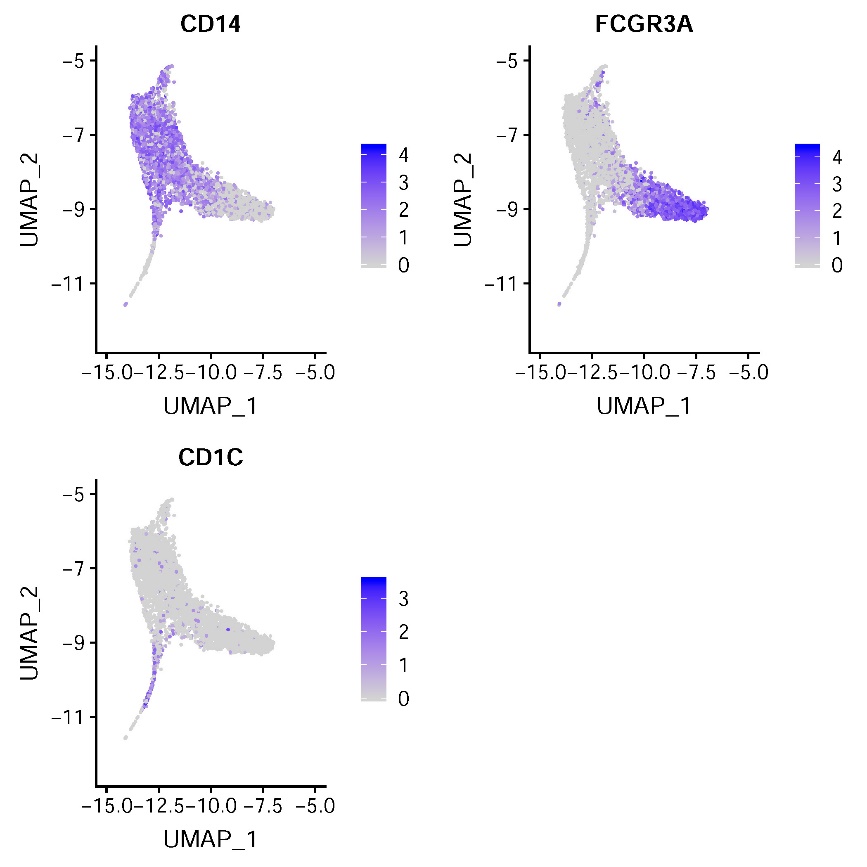

Supplementary Figure S3. Marker genes for monocyte subsets. The cells are colored based on the normalized expression of marker genes.

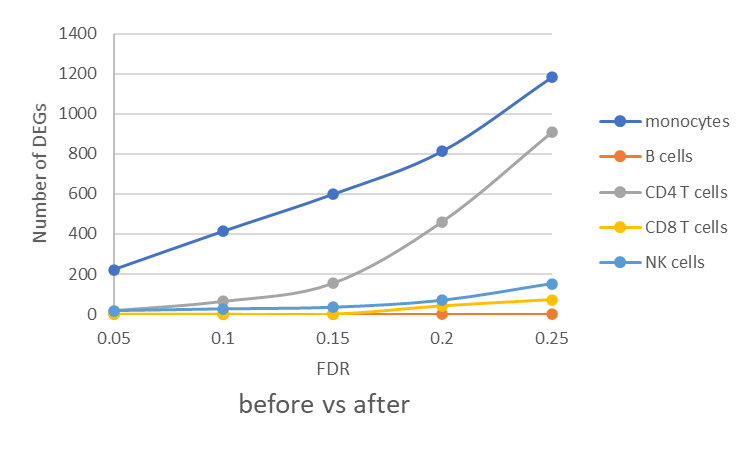

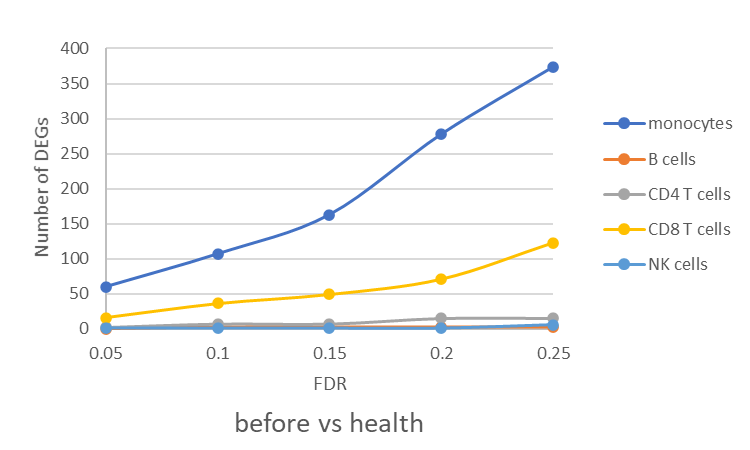

Supplementary Figure S4. Number of differentially expressed genes for each cell type. (Top) Comparison between pre- and post-therapy samples. (Bottom) Comparison between pre-therapy samples and healthy controls.

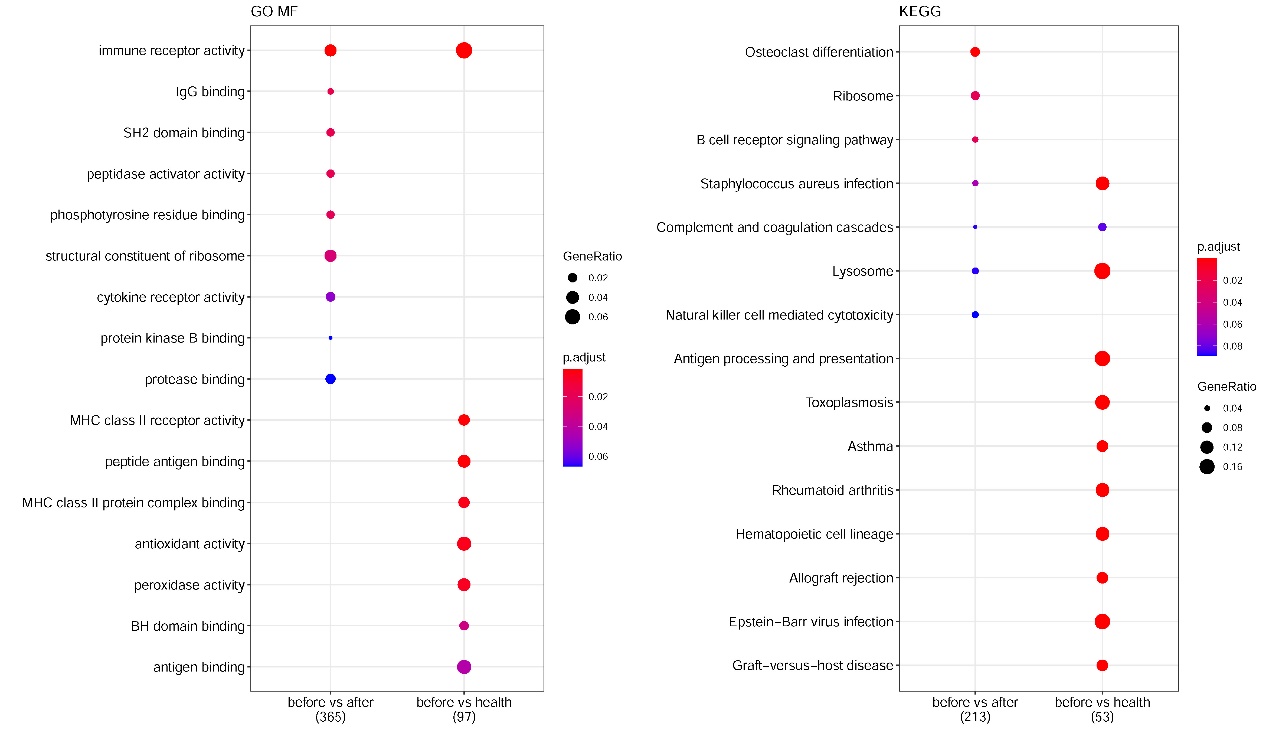

Supplementary Figure S5. Functional enrichment analyses for DEGs in monocytes. (Left) GO molecular functions. (Right) KEGG pathways. Dot size indicates gene ratio, and dot color indicates FDR.

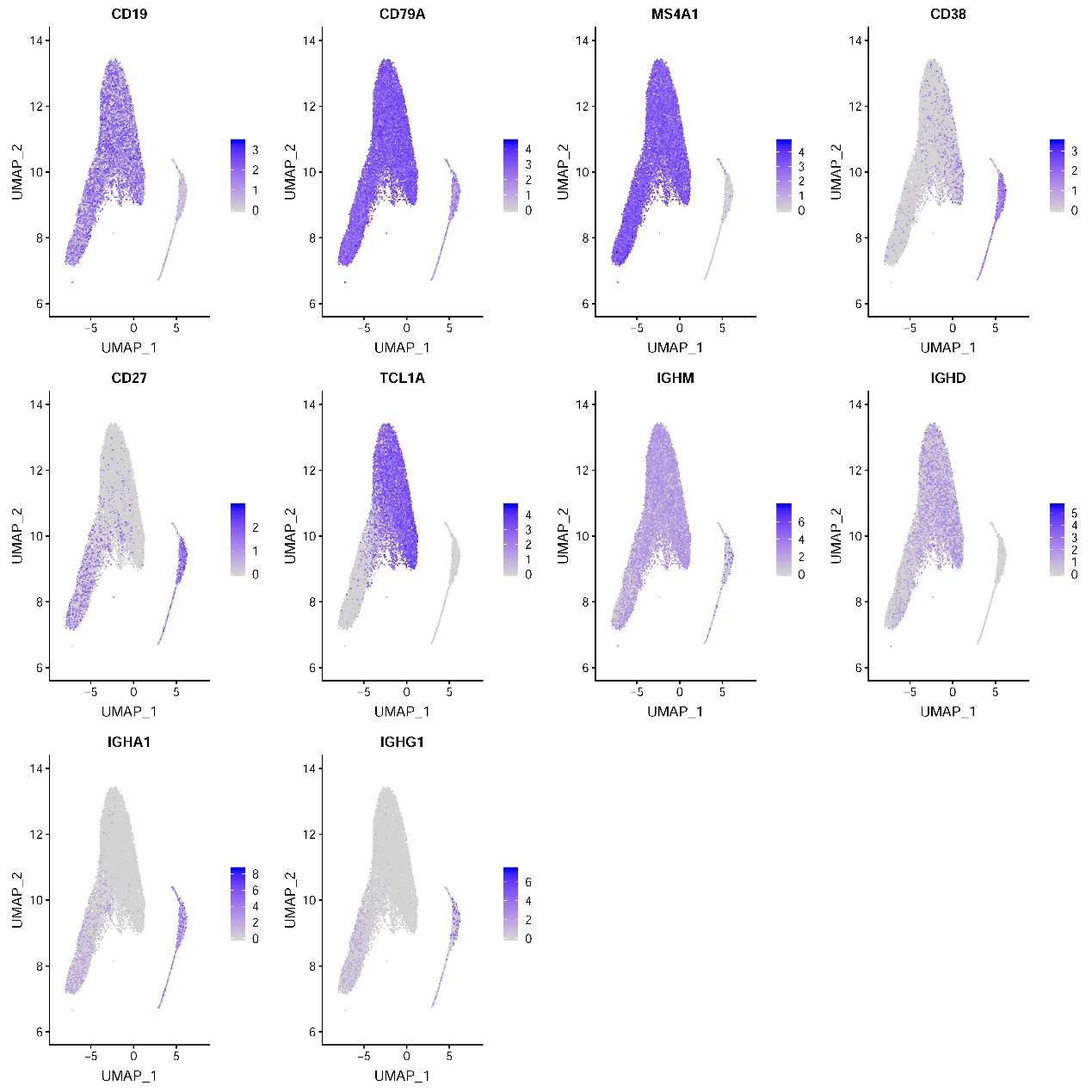

Supplementary Figure S6. Marker genes for B cell subsets. The cells are colored based on the normalized expression of marker genes.

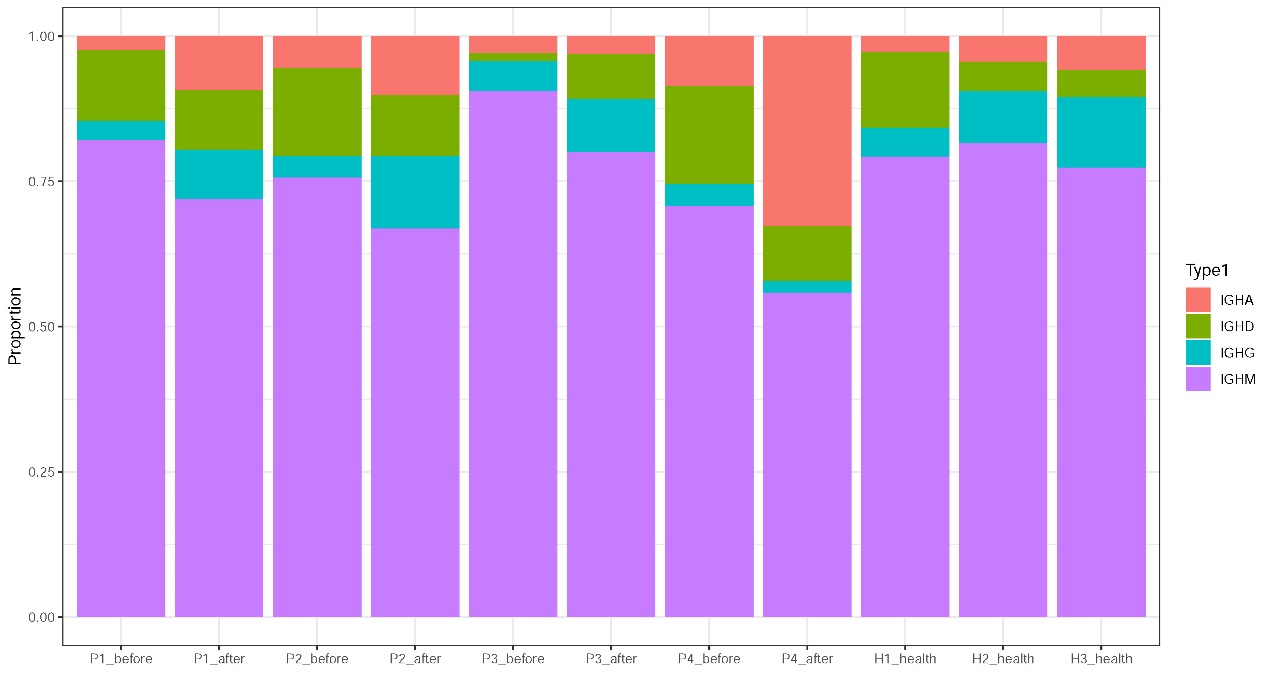

Supplementary Figure S7. Proportion of IGH isotypes in each sample.

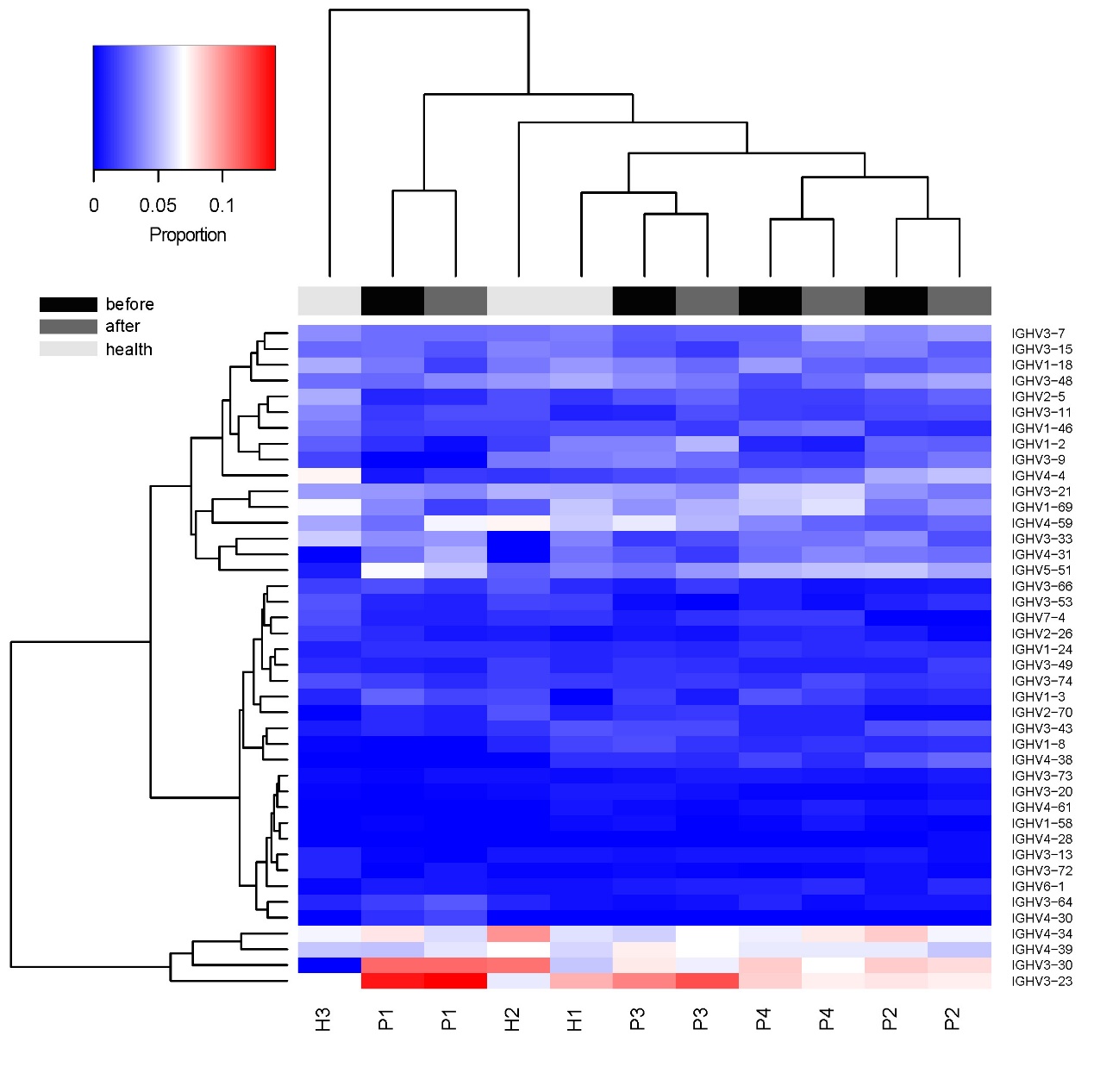

Supplementary Figure S8. IGHV gene usage in the BCR repertoires. Heat map represents proportion of IGHV genes across samples.

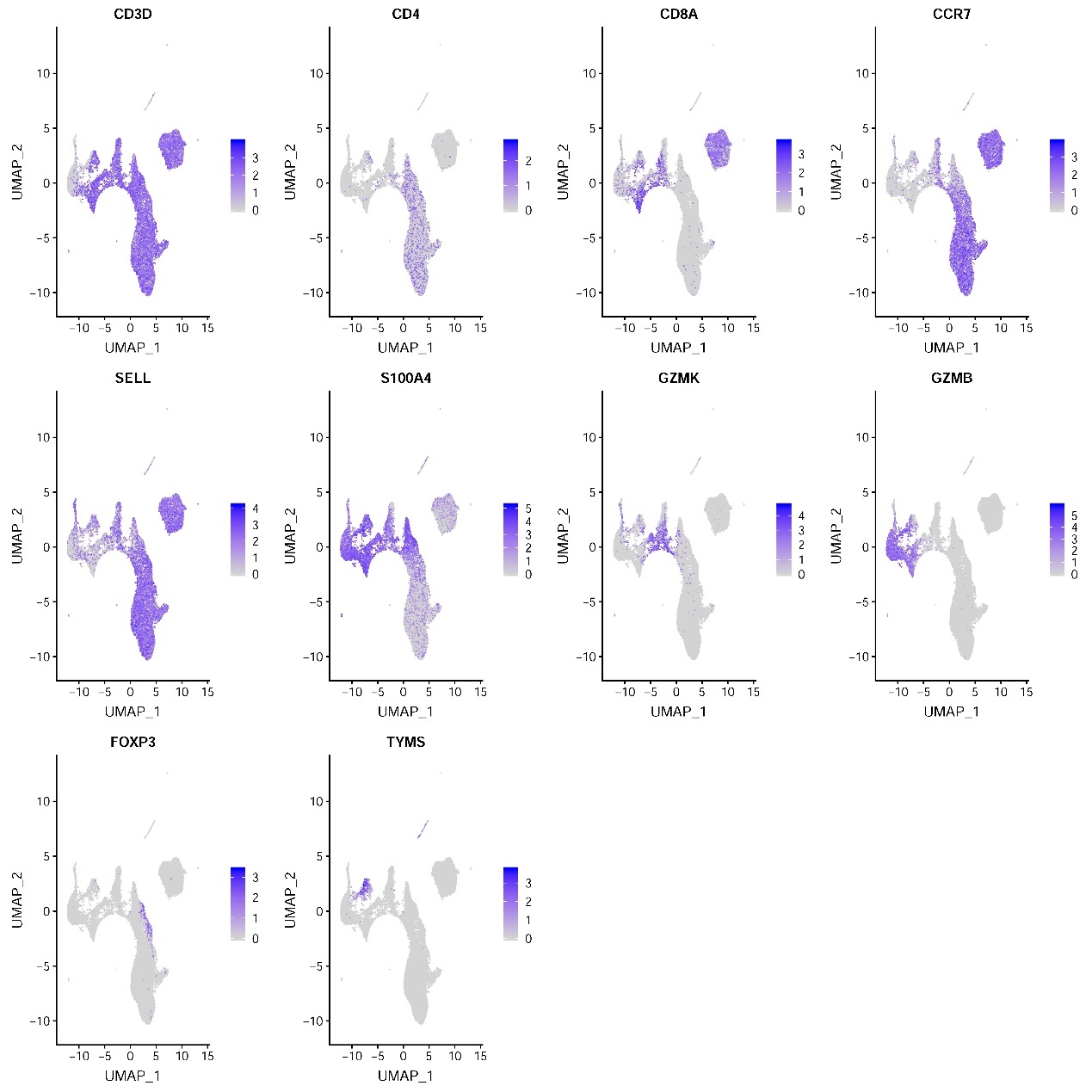

Supplementary Figure S9. Marker genes for T cell subset. The cells are colored based on the normalized expression of marker genes.

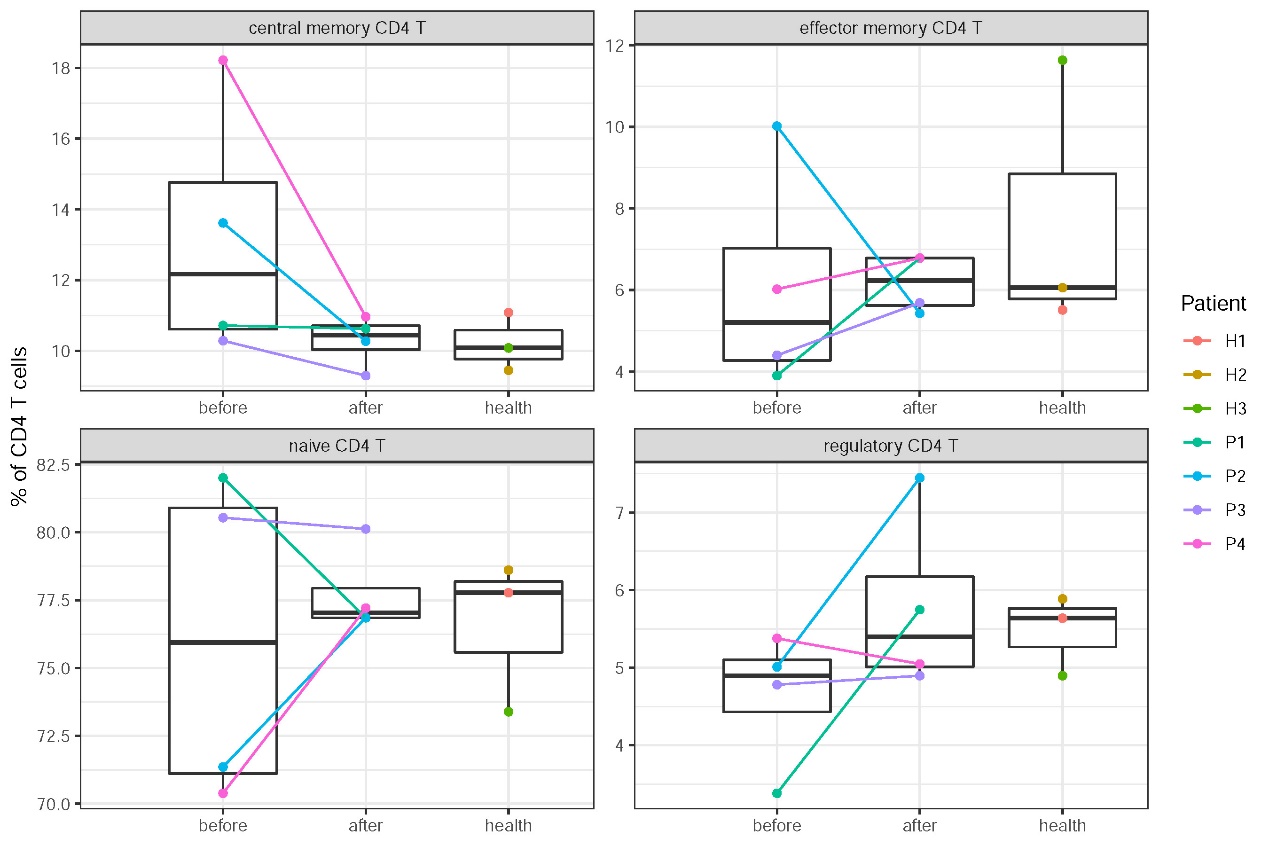

Supplementary Figure S10. Percentage of CD4 T cell subsets across conditions.

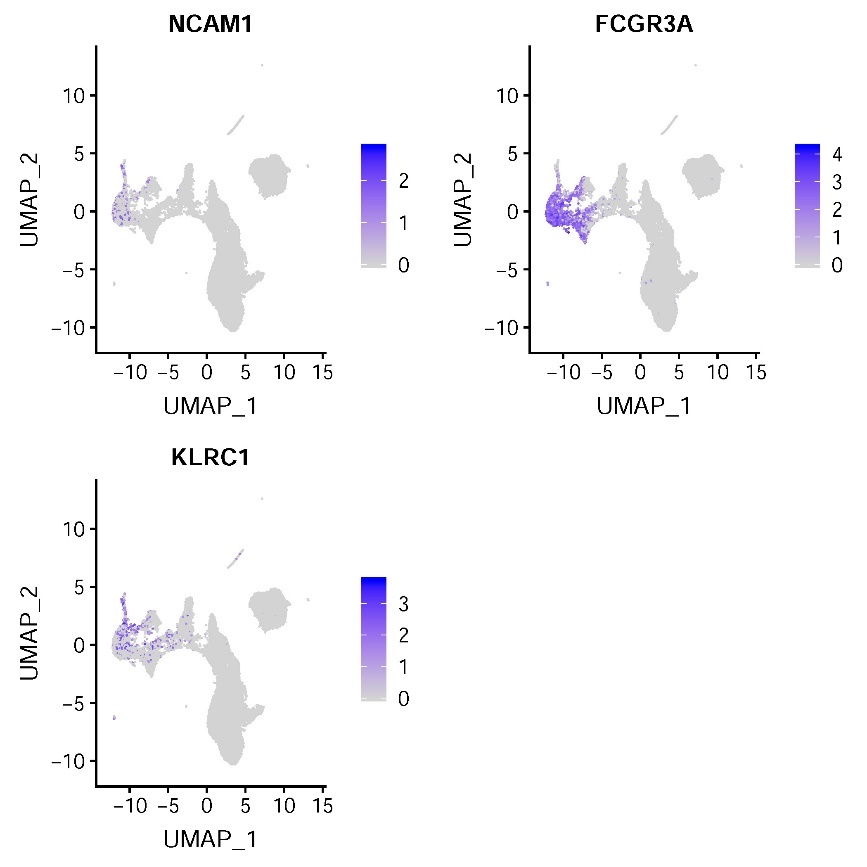

Supplementary Figure S11. Marker genes for NK cell subsets. The cells are colored based on the normalized expression of marker genes.

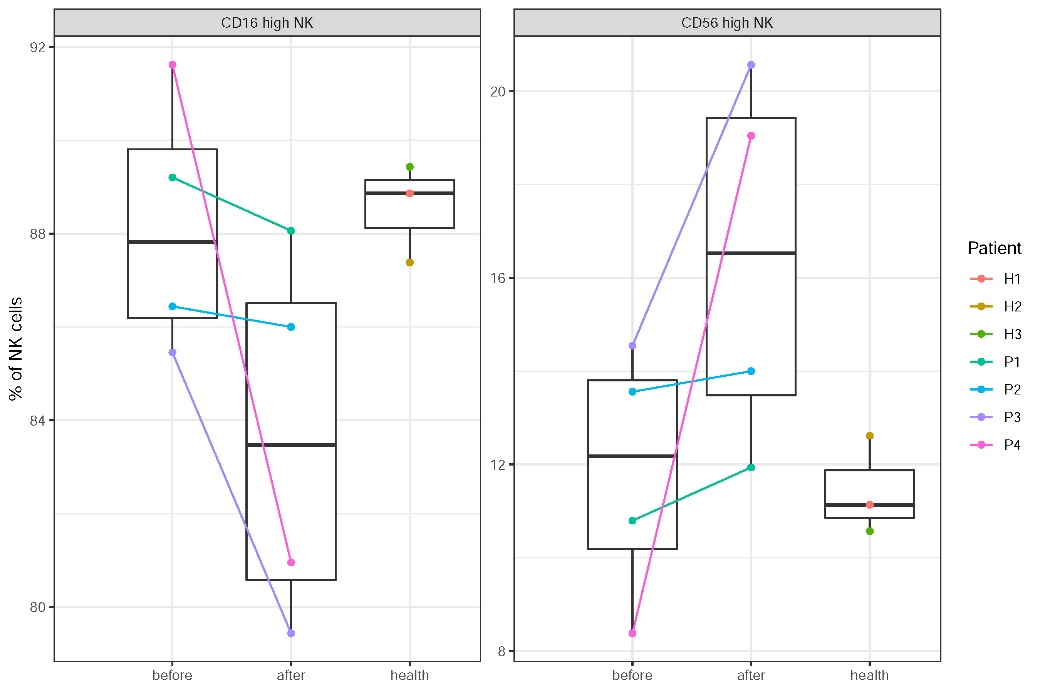

Supplementary Figure S12. Percentage of NK cell subsets across conditions.

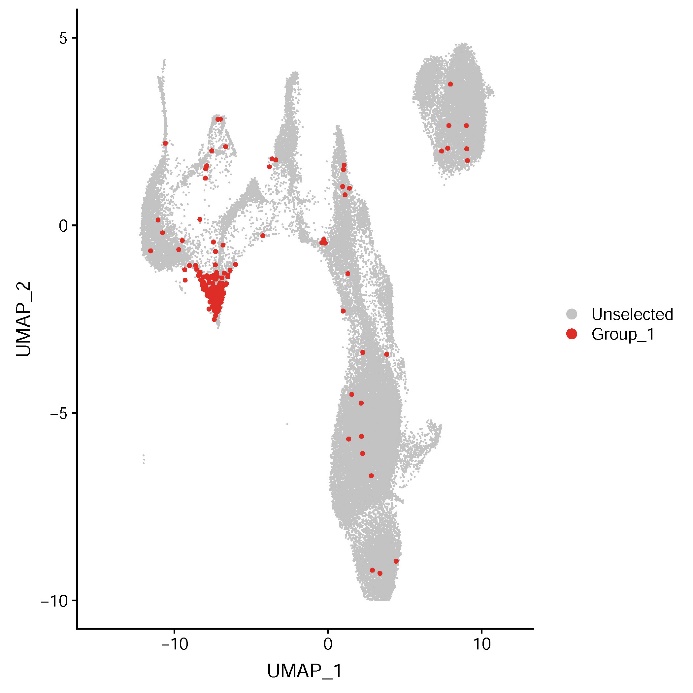

Supplementary Figure S13. T cells with clonal TCRs. T cells with clonotype size ≥ 3 are colored in red.

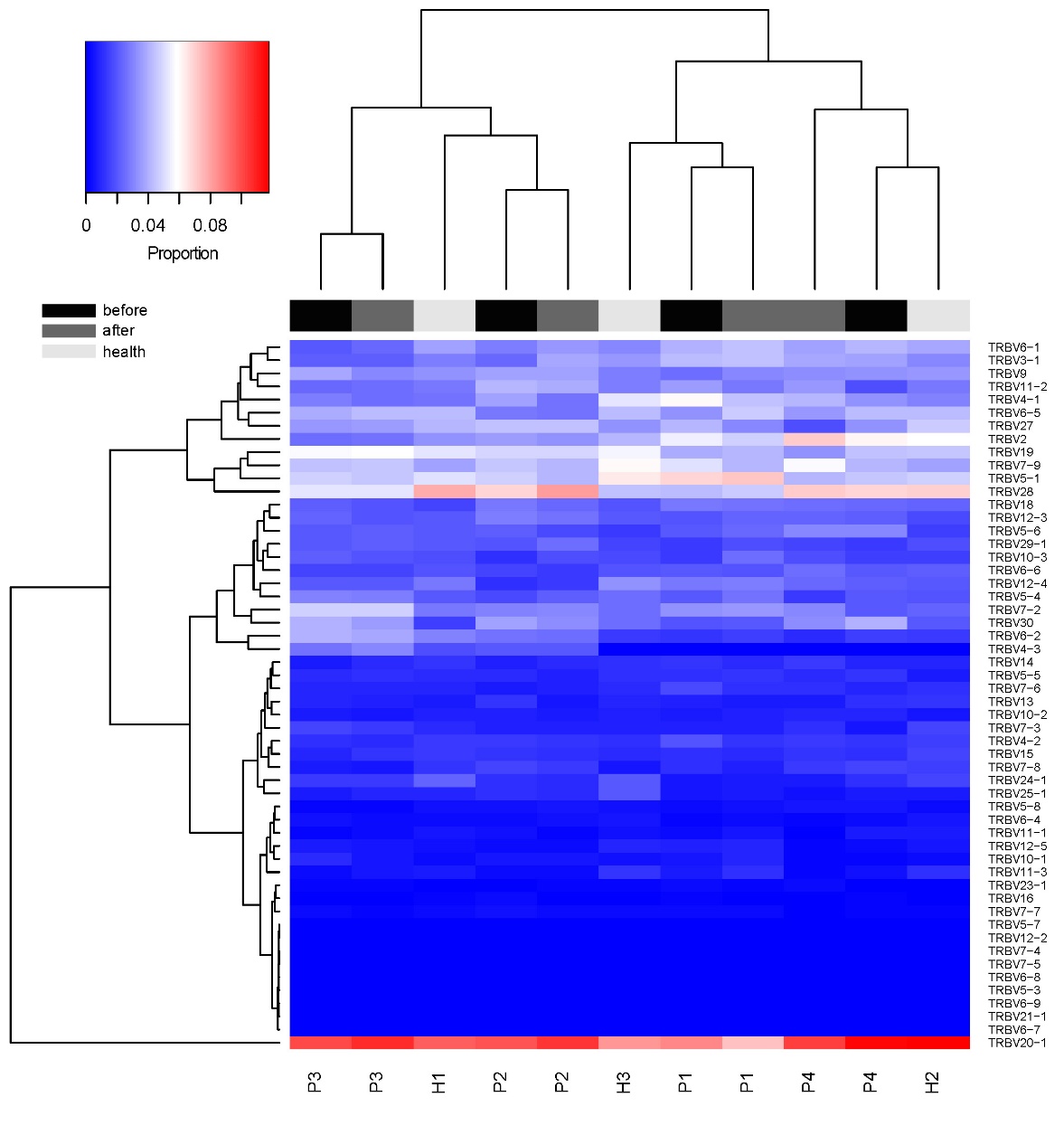

Supplementary Figure S14. TRBV gene usage in the TCR repertoires. Heat map represents proportion of TRBV genes across samples.

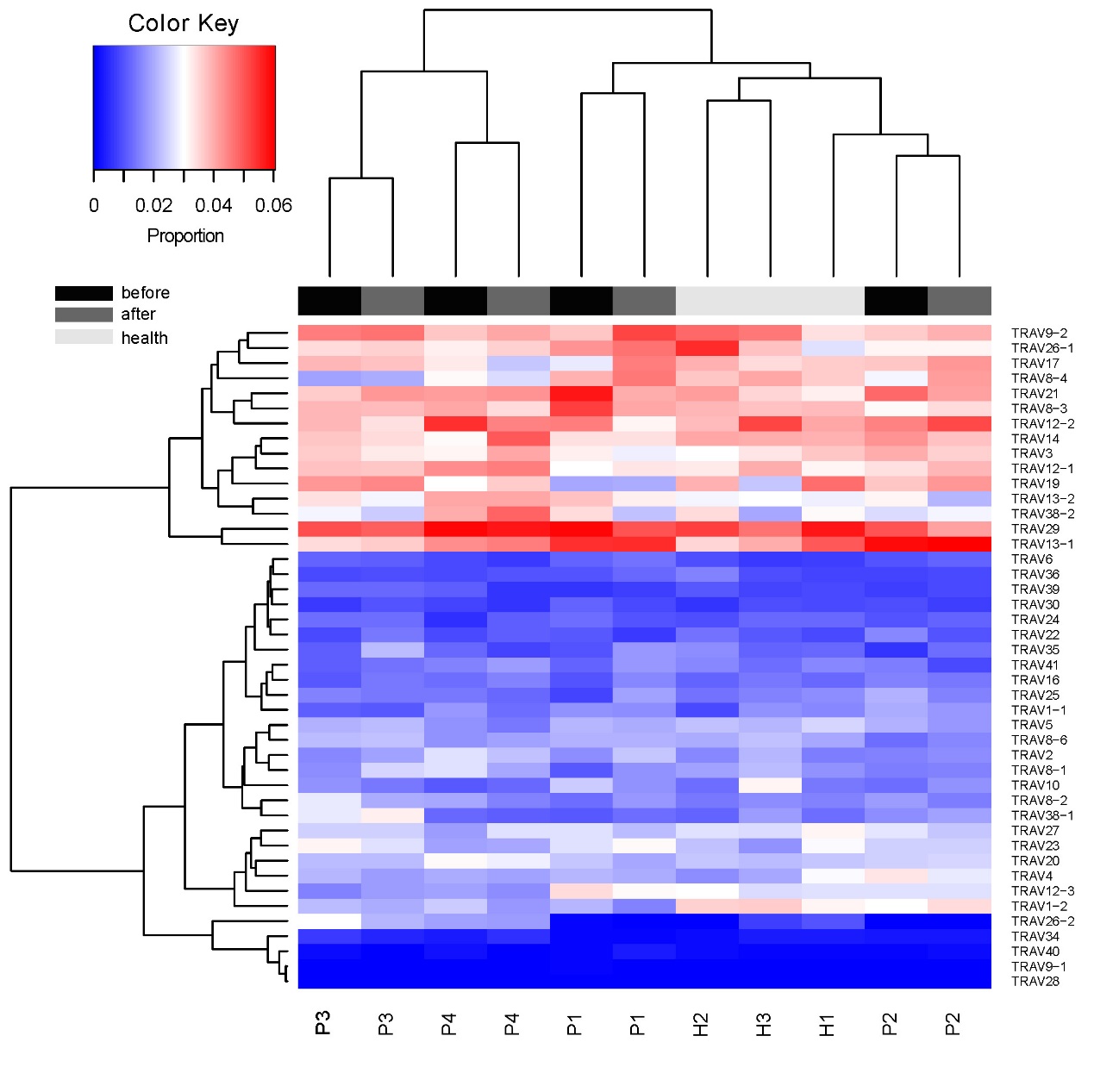

Supplementary Figure S15. TRAV gene usage in the TCR repertoires. Heat map represents proportion of TRAV genes across samples.

Supplementary Table S1. Clinical features of recruited donors.

| Donor | P1 | P2 | P3 | P4 | H1 | H2 | H3 |
| --- | --- | --- | --- | --- | --- | --- | --- |
| Age (years) | 1.6 | 5.4 | 3.3 | 2.1 | 1.7 | 3 | 5 |
| Sex (male/female) | F | F | F | M | M | F | F |
| Fever (days) | 7 | 7 | 9 | 7 |  |  |  |
| Conjunctivitis (yes/no) | Y | Y | Y | N |  |  |  |
| Oral changes (yes/no) | Y | N | Y | Y |  |  |  |
| Extremity changes (yes/no) | Y | Y | N | Y |  |  |  |
| Rash (yes/no) | Y | Y | N | Y |  |  |  |
| Cervical lymphadenopathy (yes/no) | Y | Y | Y | Y |  |  |  |
| Complete KD (yes/no) | Y | Y | N | Y |  |  |  |
| IVIG responsive (yes/no) | Y | Y | Y | Y |  |  |  |
| CAL (yes/no) | N | N | N | N |  |  |  |

Supplementary Table S2. Laboratory blood test before and after IVIG therapy.

| Patient | P1 | P1 | P2 | P2 | P3 | P3 | P4 | P4 |
| --- | --- | --- | --- | --- | --- | --- | --- | --- |
| IVIG therapy | before | after | before | after | before | after | before | after |
| White blood cell count (10^9/L) | 26.04 | 8.9 | 6.15 | 4.68 | 8.98 | 4.6 | 19.4 | 8.07 |
| Neutrophil count (10^9/L) | 19.82 | 2.79 | 4.45 | 2.98 | 5.93 | 1.72 | 14.86 | 2.22 |
| Lymphocyte count (10^9/L) | 4.04 | 3.54 | 1.45 | 1.44 | 2.38 | 2.42 | 2.99 | 4.54 |
| Monocyte count (10^9/L) | 0.57 | 1.36 | 0.23 | 0.14 | 0.49 | 0.21 | 0.33 | 0.42 |
| Red blood cell count (10^12/L) | 4.23 | 3.97 | 4.28 | 3.97 | 3.67 | 3.5 | 3.59 | 3.41 |
| Platelet count (10^9/L) | 469 | 607 | 193 | 293 | 301 | 288 | 291 | 314 |
| C-reactive protein (mg/L) | 170 | 63 | 59 | 42 | 91 | 60 | 120 | 54 |

Supplementary Table S3. Summary of single-cell RNA sequencing data.

| Donor | IVIG therapy | Number of cells | Mean reads per cell | Median UMI counts per cell | Total genes detected | Median genes per cell | Number of cells passing QC |
| --- | --- | --- | --- | --- | --- | --- | --- |
| P1 | before | 6136 | 54999 | 1480 | 17947 | 624 | 4083 |
| P1 | after | 7328 | 68558 | 4816 | 20257 | 1720 | 6042 |
| P2 | before | 6133 | 57371 | 3377 | 18898 | 1226 | 5071 |
| P2 | after | 9910 | 36008 | 4052 | 19548 | 1415 | 6885 |
| P3 | before | 6258 | 58524 | 4157 | 19094 | 1409 | 5275 |
| P3 | after | 8867 | 39405 | 5045 | 19059 | 1485 | 7640 |
| P4 | before | 5794 | 61228 | 4372 | 19627 | 1324 | 4890 |
| P4 | after | 10394 | 31982 | 1144 | 18801 | 498 | 3307 |
| H1 | health | 5697 | 65080 | 3402 | 18901 | 1263 | 5265 |
| H2 | health | 5080 | 82405 | 3265 | 18913 | 1229 | 4382 |
| H3 | health | 5471 | 59770 | 3408 | 18981 | 1336 | 5041 |

Supplementary Table S4. Summary of single-cell BCR sequencing data.

| Donor | IVIG therapy | Number of cells | Mean reads per cell | Number of cells with clonotypes | Number of clonotypes | Number of cells with productive paired chains |
| --- | --- | --- | --- | --- | --- | --- |
| P1 | before | 3242 | 9958 | 3230 | 2977 | 2546 |
| P1 | after | 904 | 27186 | 903 | 844 | 821 |
| P2 | before | 2180 | 7805 | 2170 | 2129 | 1873 |
| P2 | after | 1048 | 34832 | 1048 | 936 | 986 |
| P3 | before | 1689 | 23870 | 1685 | 1635 | 1602 |
| P3 | after | 976 | 31256 | 975 | 925 | 945 |
| P4 | before | 2695 | 10666 | 2681 | 2596 | 2290 |
| P4 | after | 7477 | 3724 | 7477 | 1102 | 954 |
| H1 | 0601 | 1525 | 21452 | 1518 | 1437 | 1446 |
| H2 | 0601 | 1313 | 15319 | 1305 | 1254 | 1231 |
| H3 | 0601 | 799 | 33742 | 792 | 780 | 725 |

Supplementary Table S5. Summary of single-cell TCR sequencing data.

| Donor | IVIG therapy | Number of cells | Mean reads per cell | Number of cells with clonotypes | Number of clonotypes | Number of cells with productive paired chains |
| --- | --- | --- | --- | --- | --- | --- |
| P1 | before | 2635 | 12020 | 2354 | 2342 | 1217 |
| P1 | after | 3810 | 6409 | 3323 | 3295 | 2912 |
| P2 | before | 3597 | 5542 | 3284 | 3162 | 2594 |
| P2 | after | 6517 | 6205 | 6352 | 5919 | 5648 |
| P3 | before | 3740 | 22339 | 3539 | 3508 | 3077 |
| P3 | after | 5028 | 14485 | 4835 | 4792 | 4400 |
| P4 | before | 3758 | 7372 | 3284 | 3273 | 2303 |
| P4 | after | 3282 | 7738 | 3187 | 3125 | 2601 |
| P5 | before | 4495 | 6799 | 4153 | 4098 | 3383 |
| P5 | after | 5519 | 6733 | 4640 | 4545 | 4120 |
| H1 | health | 4472 | 7107 | 4087 | 3658 | 3351 |
| H2 | health | 3319 | 13554 | 3016 | 2936 | 2390 |
| H3 | health | 3757 | 8035 | 3285 | 3055 | 2845 |
